# Transfer learning improves outcome predictions for ASD from gene expression in blood

**DOI:** 10.1101/2021.06.26.449864

**Authors:** Kimberly Robasky, Raphael Kim, Hong Yi, Hao Xu, Bokan Bao, Austin W.T. Chiang, Eric Courchesne, Nathan E. Lewis

## Abstract

**Background:** Predicting outcomes on human genetic studies is difficult because the number of variables (genes) is often much larger than the number of observations (human subject tissue samples). We investigated means for improving model performance on the types of under-constrained problems that are typical in human genetics, where the number of strongly correlated genes (features) may exceed 10,000, and the number of study participants (observations) may be limited to under 1,000.

**Methods:** We created ‘train’, ‘validate’ and ‘test’ datasets from 240 microarray observations from 127 subjects diagnosed with autism spectrum disorder (ASD) and 113 ‘typically developing’ (TD) subjects. We trained a neural network model (a.k.a., the ‘naive’ model) on 10,422 genes using the ‘train’ dataset, composed of 70 ASD and 65 TD subjects, and we restricted the model to one, fully-connected hidden layer to minimize the number of trainable parameters, including a dropout layer to help prevent overfitting. We experimented with alternative network architectures and tuned the hyperparameters using the ‘validate’ dataset, and performed a single, final evaluation using the holdout ‘test’ dataset. Next, we trained a neural network model using the identical architecture and identical genes to predict tissue type in GTEx data. We transferred that learning by replacing the top layer of the GTEx model with a layer to predict ASD outcome and we retrained the new layer on the ASD dataset, again using the identical 10,422 genes.

**Findings:** The ‘naive’ neural network model had AUROC=0.58 for the task of predicting ASD outcomes, which saw a statistically significant 7.8% improvement from transfer learning.

**Interpretation:** We demonstrated that neural network learning could be transferred from models trained on large RNA-Seq gene expression to a model trained on a small, microarray gene expression dataset with clinical utility for mitigating over-training on small sample sizes. Incidentally, we built a highly accurate classifier of tissue type with which to perform the transfer learning.

**Funding:** This work was supported in part by NIMH R01-MH110558 (E.C., N.E.L.)

**Author Summary:** Image recognition and natural language processing have enjoyed great success in reusing the computational efforts and data sources to overcome the problem of over-training a neural network on a limited dataset. Other domains using deep learning, including genomics and clinical applications, have been slower to benefit from transfer learning. Here we demonstrate data preparation and modeling techniques that allow genomics researchers to take advantage of transfer learning in order to increase the utility of limited clinical datasets. We show that a non-pre-trained, ‘naive’ model performance can be improved by 7.8% by transferring learning from a highly performant model trained on GTEx data to solve a similar problem.

## Introduction

Biomedicine has been slow to reap the rewards from sequencing the human genome in 2003^1^, partially because of the intractable computational complexity of permuting over the estimated 3 million variants in each human genome^2^, 30,000 genes, their RNA transcripts^3,4^, translated proteins^5^, and epigenetics. However, with modern advances in AI-driven technologies, we are closer than ever to unraveling the mysteries of the druggable genome^6^. Modern GPU servers and TPUs^7^ provide the means for delivering tremendous computational power to compute-hungry deep learning algorithms that can capture all the nonlinear and multivariate correlations between longitudinal clinical covariates and biomolecular data^8,9^. However, to learn the statistical relationships between so many features, these algorithms require vast amounts of data and human biosamples are limited while data access is constrained by processes aimed at protecting patient privacy and confidentiality.

Fortunately, emerging AI methods born from image recognition research^10^ are driving advances in biomedical research that are beginning to translate into clinical applications, such as radiology^11^ and histopathology^12^. Transfer learning^13,14^ is one of the breakthroughs empowering these advances. Now, a model developer can transfer the learning from a model trained on a very large dataset to solve a similar problem with a new model trained on a relatively smaller dataset. This approach saves on data collection costs and time as well as reduces the number of patients to recruit and consent for study. However, transfer learning has largely only been useful for image-based problems that lend themselves to convolutional neural networks (CNNs)^15^, with some emerging applications in natural language processing (NLP)^16^, and in omics^17^. CNNs are complicated and still require a significant amount of data, but what if we could pre-train a simple, multilayer perceptron (MLP)^18^ network for use on even smaller datasets?

Given access to a fully de-identified, limited dataset of human subjects, we tested if transfer learning of a simple neural network trained from a large public database of well-annotated gene expression could potentially be used for developing an ASD classifier for decision support in the clinic. Armed with gene expression microarray data and binary clinical outcomes of a mere 240 individuals, we successfully applied transfer learning to improve performance of a very simple MLP for classifying 12-18 month old pediatric patients as either TD or ASD from a simple blood sample.

## Methods

### Study population datasets

This cohort is described elsewhere (manuscript in preparation) and summarized here: Leukocyte-based transcriptomics were collected on male toddlers ages 1 to 4 years. More than half were recruited from the general population as young as 12 months using an early screening, detection, and diagnosis strategy called the Get SET Early procedure^19^ and the remaining were from general community referrals and the final diagnostic evaluation occurred between the ages of 18 to 24 months. The toddlers studied here were among those in a >1,200 sample study of early diagnostic stability in ASD^20^. Transcriptome data based on microarray methods (see below) were previously published on (147 individuals)^20,21^. The Institutional Review Board of the University of California, San Diego, approved research procedures. Parents of subjects underwent Informed Consent Procedures with a psychologist or study coordinator at the time of their child’s enrollment. Four to six milliliters of blood were collected into ethylenediaminetetraacetic-coated tubes from all toddlers. Blood leukocytes were captured and stabilized by LeukoLOCK filters (Ambion) and were immediately placed in a −20°C freezer. Total RNA was extracted following standard procedures and manufacturer’s instructions (Ambion). Gene expression was assayed using Illumina HumanHT-12 Expression beadchip. All arrays were scanned with the Illumina BeadArray Reader and read into Illumina GenomeStudio software (version 1.1.1). Raw Illumina probe intensities were converted to expression values using the Lumi Bioconductor package^22^ and probes were filtered for reliable expression levels culminating in the selection of 14,312 expressing coding genes.

For this study, we subsampled 240 male individuals composed of 127 toddlers with ASD diagnoses and 113 typically developing (TD) toddlers. Previous efforts uncovered analytical difficulties with classification of females in this domain, and with our goal of exploration of modelling methods, we deemed it unnecessary to gender-balance the data.

We downloaded GTEx gene expression data from https://greenelab.github.io/BioBombe/^23^, retaining the author’s training and validation splits. We retained only RNA-Seq samples (e.g., where column ‘SMAFRZE’==’RNASEQ’), dropping 3,910 samples. Data are more fully summarized in Table 1.

**Table 1.**
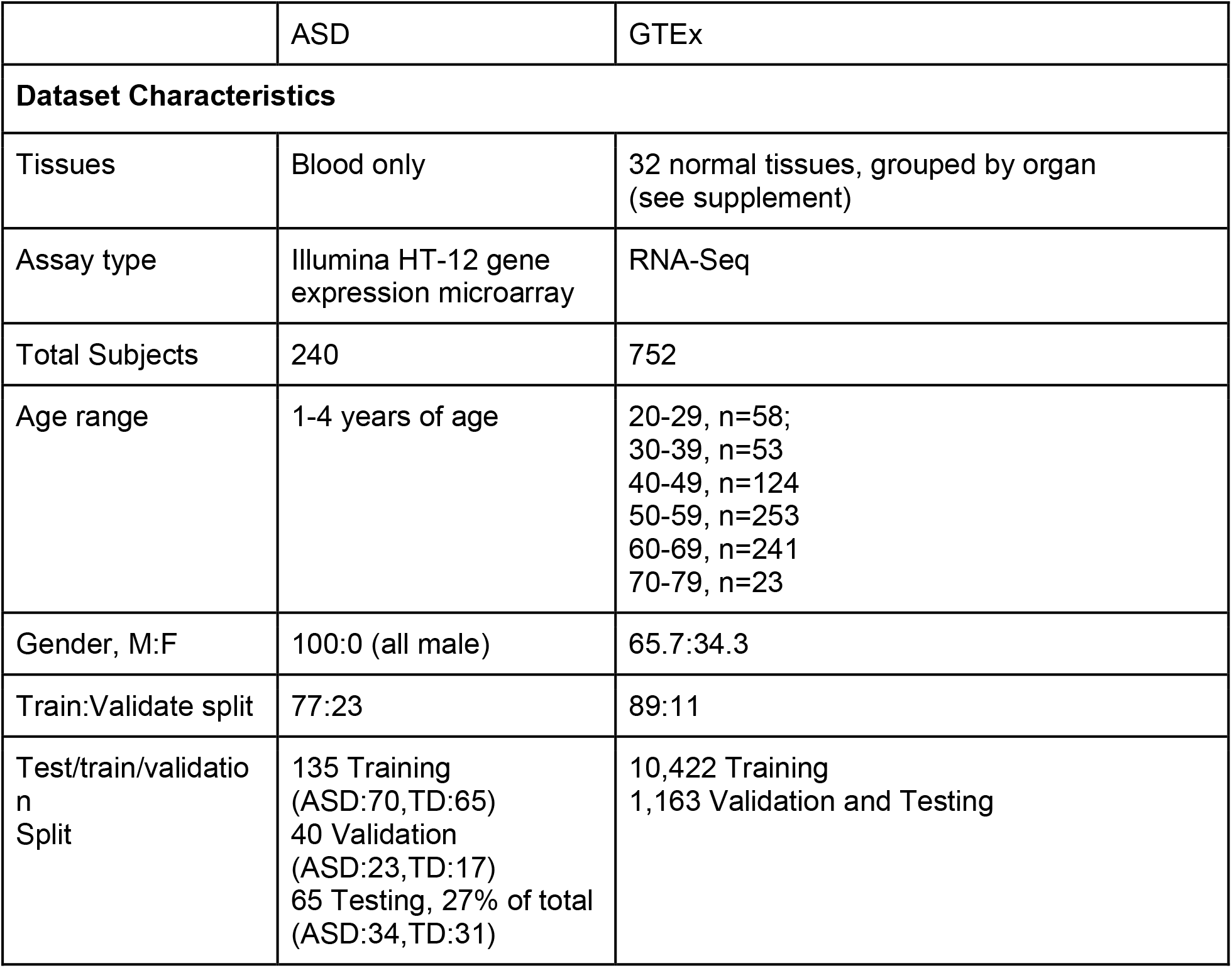
Summary of the training, validation, and test dataset characteristics.

|  | ASD | GTEx |
| --- | --- | --- |
| **Dataset Characteristics** |  |  |
| Tissues | Blood only | 32 normal tissues, grouped by organ (see supplement) |
| Assay type | Illumina HT-12 gene expression microarray | RNA-Seq |
| Total Subjects | 240 | 752 |
| Age range | 1-4 years of age | 20-29, n=58;<br>30-39, n=53<br>40-49, n=124<br>50-59, n=253<br>60-69, n=241<br>70-79, n=23 |
| Gender, M:F | 100:0 (all male) | 65.7:34.3 |
| Train:Validate split | 77:23 | 89:11 |
| Test/train/validation Split | 135 Training (ASD:70,TD:65)<br>40 Validation (ASD:23,TD:17)<br>65 Testing, 27% of total (ASD:34,TD:31) | 10,422 Training<br>1,163 Validation and Testing |

Feature extraction and dataset splitting GTEx sample types were grouped by organ such that all brain tissue subtypes were labeled as brain, all kidney tissue subtypes labeled as kidney, and so forth. This resulted in reducing the tissue type classes from 54 to 32. See supplemental table S1 for sample counts per tissue type.

From our 240 samples of ASD data, a holdout test dataset of 65 was selected by our data provider colleagues, leaving 175 samples for our discovery set. We then randomly split our discovery set into training (*n=135*) and validation (*n=40*). Due to the limited size of the dataset, we did not under-sample for balance between TD and ASD, which we determined was sufficient for comparing performance between models. If we were aiming to optimize performance, e.g., for a diagnostic test, we would analyze on the larger dataset, currently in preparation, and we would subsample to exactly balance the number of ASD and TD in the training.

### Model development

We trained a neural network classifier to predict ASD and TD outcomes. We used a simple, multilayer perceptron (MLP) neural network with a single hidden layer whose nodes are equal to the average of the input and ASD output layers (5,674). We used 0.50 dropout rate in our dropout layer to accommodate under constrained variables helping to prevent overfitting. (Figure 1, a.)

**Figure 1.**
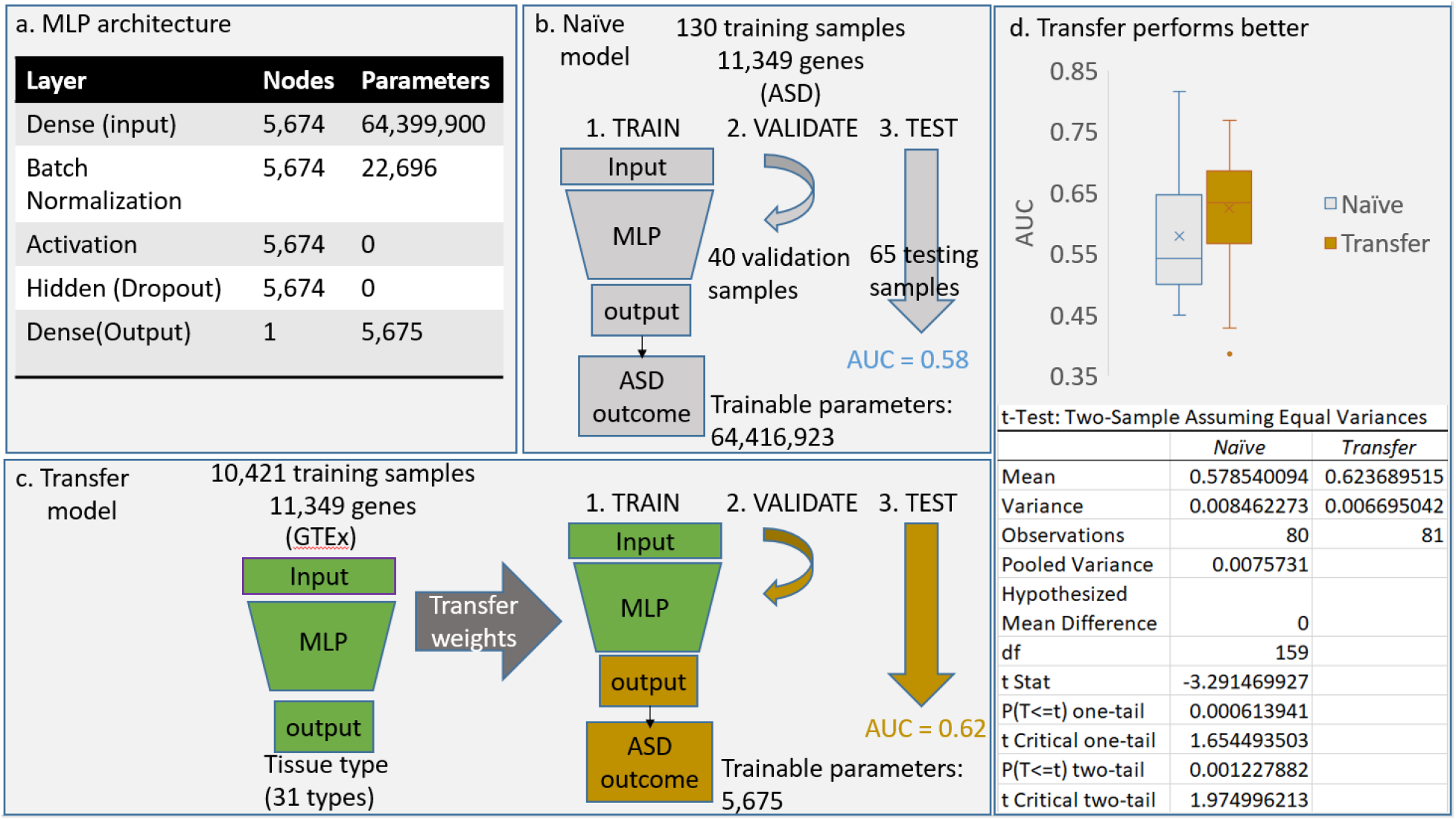
Training, evaluation pipeline and results. **a.**We used an identical multilayer perceptron (MLP) neural network architecture with a single hidden layer for each experiment. **b.**We trained the 64,416,923 parameters for the ‘naive’ model on 130 samples with 11,349 gene features and a binary ASD outcome target. We chose the MLP architecture and hyperparameters using 40 validation samples and we used the keras API to compute the mean AUC=0.58 on 80 training rounds, evaluating each round with a hold-out test dataset of 65 samples. **c.**We trained the same architecture on 10,421 GTEx samples to predict 1 of 31 tissue types. We then transferred those weights, locking all but the output layer, resulting in 5,675 trainable parameters, and repeated the experiment described in b. We found that the mean AUC=0.62 **d.**We performed a t-Test on the AUC distributions, α=0.05; the difference between the means is statistically significant.

### Training

Given the limited sample size and overwhelming number of trainable parameters, we used the validation data set to select the best number of epochs to use for avoiding overfitting. We tried a titration style approach where we trained under 50, 100, 200, 300, and 500 epochs to optimize the ‘area under the receiver operator curve’ (AUROC, a.k.a., AUC)^24^. We also tried an orthogonal method, ‘early stopping’, implemented with the keras API. For the keras early stopping method, we set the ‘patience’ to a higher value for the under-constrained ASD model than for the GTEx model. In both the titration and early stopping methods, we found 100-200 epochs optimized the mean AUC across multiple training rounds on the validation dataset, with 200 yielding more stable AUC results. For repeatability, we chose to move forward to evaluation using a fixed number of epochs, 200. Other parameters used to train the models are described in Table 2 and the training approach is illustrated in Figure 1, b.

**Table 2.** Training parameters. used on each model.

| parameter | ASD, naive | ASD, pre-trained | GTEx |
| --- | --- | --- | --- |
| Learning rate(LR) | 0.001 | 0.001 | 0.001 |
| Patience | 20 | 20 | 10 |
| LR Patience | 2 | 2 | 2 |
| Early stopping/Validation samples | 30/10 | 30/10 | 30/10 |
| epochs | 200 | 200 | n/a (early stopping) |

### Pre-training

We aimed to optimize trainable parameters by transferring the weights from an MLP with more observations that can solve a similar problem. For this we choose the GTEx dataset^25^, which has a more substantial number of observations (11,585), that can be used to solve the similar problem of tissue classification. Because the target dimension differs from task to task (e.g. *n*_1_*∊* {0, 1, …,31} denoting tissue types in GTEx and *n*_2_*∊* {0, 1}denoting outcomes in our ASD dataset), when transitioning from one training task to the next, the last layer is removed and re-initialized randomly to a new layer with the proper size for the target space. Refer to Figure 1, c. for a depiction of the pre-training process. Since the aim of this experiment was not to optimize the GTEx model but rather to analyze any relative differences made by pre-training, we did not tune hyperparameters for the GTEx model beyond implementing the keras API’s early stopping for selecting the number of epochs. We used a small 40-sample holdout from the large 1,163-sample test dataset for the early stopping method, indicated in Table 3. We used the full GTEx test dataset (1,163 samples) for evaluating that model. We used 40 samples for the ASD models to evaluate hyperparameters.

**Table 3.**
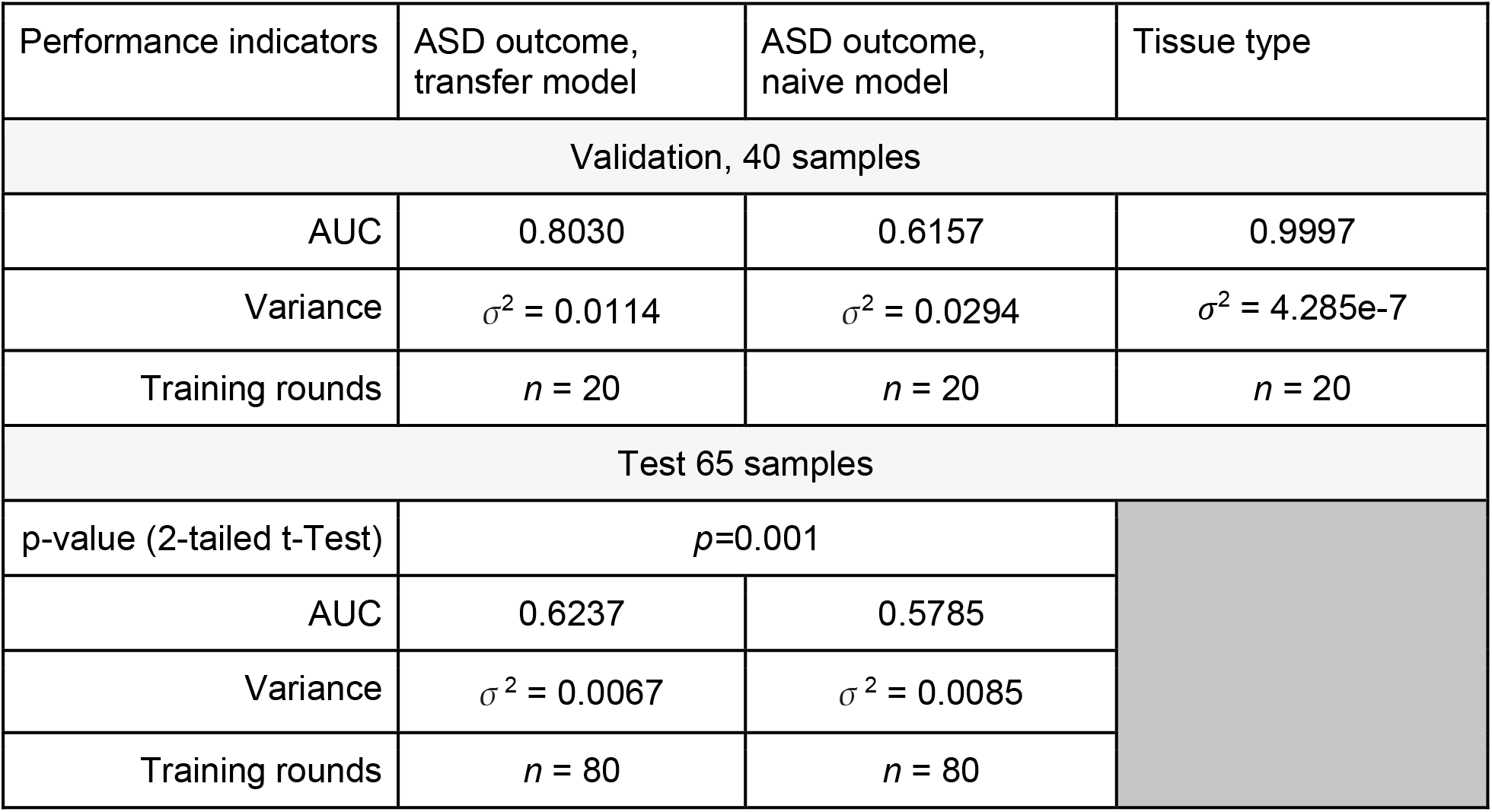
Classification performance.

| Performance indicators | ASD outcome,<br>transfer model | ASD outcome,<br>naive model | Tissue type |
| --- | --- | --- | --- |
| Validation, 40 samples |  |  |  |
| AUC | 0.8030 | 0.6157 | 0.9997 |
| Variance | $\sigma^2 = 0.0114$ | $\sigma^2 = 0.0294$ | $\sigma^2 = 4.285e-7$ |
| Training rounds | $n = 20$ | $n = 20$ | $n = 20$ |
| Test 65 samples |  |  |  |
| p-value (2-tailed t-Test) | $p=0.001$ | | |
| AUC | 0.6237 | 0.5785 |  |
| Variance | $\sigma^2 = 0.0067$ | $\sigma^2 = 0.0085$ | |
| Training rounds | $n = 80$ | $n = 80$ | |

### Code and training platform

We trained the models on a standalone graphical processing unit (GPU) server with two NVidia Tesla P100 cards and 755GB RAM. Code and models were read from a network attached storage (NAS) device and performance on these hardware devices are reported (Table 3). All code and supplemental data are publicly available and version controlled at https://github.com/RENCI/tx-GeneXNet).

## Results

The test AUC performance measurement saw a 7.5% improvement when using transfer learning (see Figure 1, d.) over the non-pre-trained model. A two-tailed T-test confirmed the statistical significance of the difference in the average AUC over 60 training rounds of the pre-trained model, versus the model that was not pre-trained (see Table 3).

The validation dataset, which was used to tune hyperparameters, out-performs AUC for holdout test dataset (see Table 3). Variance for the test dataset, which had 80 rounds of training, is smaller than for the validation dataset, which had only 20 rounds. There is a small (*Δ*=0.0452), but statistically significant (*p=*0.001) difference between naive and transfer models. Notably, the pre-trained model trained on tissue type outcomes performs near optimally with an AUC of .9997 on unseen data and very low variance.

Computational costs were relatively similar for training with both models, and considerably higher for the model trained on the GTEx model (Table 4).

**Table 4.** Compute costs.

|  | ASD outcome,<br>transfer | ASD outcome,<br>naive | Tissue type,<br>naive |
| --- | --- | --- | --- |
| training rounds,<br>epochs | 40 rounds,<br>200 epochs | 40 rounds,<br>200 epochs | 20 rounds<br>Early stopping:<br>~22 epochs |
| Real | 31m<br>41.247s | 31m<br>18.771s | 127m<br>29.969s |
| User | 61m<br>53.351s | 61m<br>30.214s | 250m<br>35.840s |
| Sys | 11m<br>3.594s | 10m<br>43.889s | 49m<br>51.713s |

## Discussion

Even the most modest performance improvements in diagnostics have clinical utility, and this study demonstrated that transfer learning, even on a simple MLP with non-image data, can achieve that aim. The average AUC of the pre-trained model is relatively low, but still impressive given an experiment with 10,422 variables and only 240 total observations. This approach additionally demonstrates a method for combining RNA-Seq data with gene expression microarray data. Other studies have transferred learning from images, and even images of gene expression, but transferring directly from counts is new territory.

Furthermore, the highly performant (AUC=1.0) tissue type classifier trained in this work has additional utility to inform primary site for cancers of unknown origin (CUP) and as an automated sample validation step in bioinformatics pipelines. An open problem in oncology practice is determining therapy for metastatic cancers where the original cancer site was not determined, and a tissue type classifier can inform where to look for the primary tumor. More research is needed to inspect the utility of this approach on tumor-specific assets like TCGA. This kind of tissue classifier can also aid in detection of sample validation switches in labs that receive biosamples externally from multiple research and clinical sources. Sample switches are rare, but highly problematic, and commercial sequencing pipelines can and do include organism detection algorithms, so in those cases, including a tissue type classifier would be a simple, low-cost measure for including one more quality control step.

Next, we aim to leverage these findings and others to optimize the performance of a diagnostic blood test for informing interventions how to help children at risk for ASD. Optimization efforts will include analyzing a larger dataset, reducing the feature set, and integrating other predictive technologies. Efforts are already underway to sequence and annotate even more samples that are being generated by our collaborators. We also aim to optimize modeling on this larger dataset by designing the output layer to compute a risk score and combining it with clinical covariates. We will tune for specificity over sensitivity to maximize clinical utility, given that false positives are particularly pernicious in this context. Finally, we aim to build in prior knowledge about gene relationships via an adjacency matrix using Graph Convolutional Networks (GCNs). We will combine these activities with active learning^26^ to enable continuous improvement as more ASD domain gene expression datasets and clinical covariates are consented for research in the community.

Transfer learning has revolutionized image recognition technology, resulting in disruptive changes to manufacturing, automobile, and medical imaging industries, and we imagine a similar future for transfer learning in gene expression analysis. We hope that these activities will lead to the sharing of pre-trained models of gene expression classification, similar to MNIST^27^ CIFAR^28^, and ImageNet^29^. With such public tools, sourced ethically and openly, we anticipate exciting biomolecular advances in this postgenomic era.

